# Myxosortases process MYXO-CTERM and other bacterial C-terminal protein-sorting signals that have invariant Cys residues

**DOI:** 10.1101/2023.01.19.524832

**Authors:** Daniel H Haft

## Abstract

The LPXTG protein-sorting signal, found in surface proteins of various Gram-positive pathogens, was the founding member of a growing panel of prokaryotic small C-terminal sorting domains. Sortase A (SrtA) cleaves LPXTG, exosortases (XrtA and XrtB) cleave the PEP-CTERM sorting signal, archaeosortase A (ArtA) cleaves PGF-CTERM, and rhombosortase (RrtA) cleaves GlyGly-CTERM domains. Three sorting signal domains without previously known processing proteases are the MYXO-CTERM, JDVT-CTERM, and SYNERG-CTERM domains. These exhibit the standard tripartite architecture of short signature motif, then a hydrophobic transmembrane segment, then an Arg-rich cluster. Each has an invariant cysteine in its signature motif. Here, we show computational evidence that these three Cys-containing sorting signals are processed by corresponding subfamilies of glutamic-type intramembrane proteases, related to type II CAAX-processing proteases found in eukaryotes. We name these sorting enzymes generally as myxosortases, and identify MXAN_2755 from Myxococcus xanthus as MrtX (myxosortase X). Additional myxosortases families MrtC and MrtP have radically different N-terminal domains, suggesting most myxosortases act as bifunctional enzymes. Myxosortase-like processing enzymes are identified also for the JDVT-CTERM (MrtJ) and SYNERG-CTERM (MrtS). This work establishes a major new family of protein-sorting housekeeping enzymes for the surface attachment of proteins on bacterial outer membranes.

## Introduction

In previous bioinformatics investigations, we identified a number of short, C-terminal protein-sorting signals in bacteria and archaea, then identified their respective processing enzymes as distinct novel proteases with multiple membrane-spanning alpha helices and with active site residues inside or near the surface of the plasma membrane. As a rule, proteins bearing these sorting signals are known or expected to undergo cleavage that removes C-terminal sequence, leaving the mature form of the target protein anchored covalently to the cell surface. That previous work focused on sorting signals that share a standard tripartite pattern of design with the classical LPXTG motifs of target proteins in Gram-positive species such as *Staphylococcus aureus*(1) and *Streptococcus pneumoniae*(2) plus sequence from the motif to the C-terminus. The three parts of the overall sorting signal are 1) a signature motif, 2) a hydrophobic segment appropriate in length for a transmembrane alpha-helix, and 3) a cluster of basic amino acids, usually several Arg residues, at or close to the protein C-terminus. We used analogy to the prototypical system that pairs LPXTG sorting signals (1) with the sortase enzyme able to process them (3), in the absence of any recognizable homologies to any parts of that system, to drive discovery and interpretation of multiple novel sorting systems(4).

The PEP-CTERM sorting signal was the first we found purely through *in silico* analysis (5). As with LPXTG proteins, we frequently observed 20 or more proteins per proteome bearing this C-terminal region, all appearing to have N-terminal signal peptides as well. Within any one genome studied, most PEP-CTERM proteins lacked any other regions of sequence similarity to any other PEP-CTERM proteins. The system was found to be more widespread than LPXTG systems, but less conspicuous because of its absence from known bacteria pathogens. PEP-CTERM systems are sporadically distributed, in Proteobacteria, Cyanobacteria, and multiple other lineages of bacteria that have a periplasm and an outer membrane. The putative sorting enzyme, a highly hydrophobic protein with eight putative transmembrane alpha-helices, which we identified by strict co-occurrence with PEP-CTERM across a large number of genomes, frequently is found within EPS (extracellular polysaccharide, or extracellular polymeric substance) biosynthetic loci. For that reason, this deeply membrane-embedded putative processing protein for PEP-CTERM proteins was named exosortase. We proposed that PEP-CTERM/exosortase systems contribute to biofilm and floc formation by large numbers of environmental organisms.

Supporting experimental work has since shown that disrupting expression of PEP-CTERM proteins disrupts floc formation in *Zoogloea resiniphila*, isolated from an activated sludge wastewater treatment plant(6). Reintroduction of the PEP-CTERM protein PepA on a plasmid restores floc formation. These findings fit with observations that most PEP-CTERM proteins lack homology to known families of enzymes and that many have low-complexity regions rich in Thr and Ser residues, suggesting extensive glycosylation. The direct demonstration that PEP-CTERM proteins are required for flocculent rather than planktonic growth further supports a model of protein anchoring on the cell surface, rather than release into the extracellular milieu, and thus further extends the analogy to the LPXTG/sortase system.

Homologs to (bacterial) exosortases occur in a number of archaeal halophiles and archaeal methanogens, and are called archaeosortases(7). The PGF-CTERM sorting domain occurs at the C-terminus of the S-layer-forming major cell surface glycoprotein of *Haloferax volcanii*. In that species, the archaeosortase ArtA is required for two linked (possibly simultaneous) processes, removal of the C-terminal alpha-helix that is part of the PGF-CTERM domain, and attachment of a large prenyl-derived lipid that sits in the membrane(8,9). Patterns of amino acid conservation in multiple sequence alignments, and site-directed mutagenesis studies of *artA* suggested by thos patterns, both support identification of exosortases and archaeosortases as novel cysteine proteases from a previously unrecognized protease family. It is not yet clear whether or not archaeosortase is a transpeptidase that removes the original protein C-terminus and replaces it with a large lipid moiety in a single step. Transpeptidation can be suspected because sortase A, an unrelated protein with a similar Cys, Arg, and His catalytic triad, performs one-step transpeptidations on LPXTG-CTERM proteins, leaving the target proteins shorter at the C-terminus and attached covalently to the Gram-positive cell wall (1,3). The target protein, transiently attached to SrtA’s active site Cys residue after the initial cleavage, is transferred from there to a cell wall precursor molecule (transpeptidation), rather than to water (hydrolysis).

Both exosortases and archaeosortases have multiple distinctive subfamilies that act, apparently, on distinct and often readily separated sets of target proteins that are marked by different flavors of sorting signal(7). However, not all sorting signals we discovered could be paired to an archaeosortase or exosortase. The GlyGly-CTERM system was one notable exception. It strictly co-occurs with (and thus likely is processed by) rhombosortase, a member of the rhomboid family of intramembrane serine proteases(10). In 2018, Gadwal, *et al*. (11) experimentally confirmed our *in silico* identification of rhombosortase in *Vibrio cholerae*. They furthermore placed the cleavage site at the C-terminal side of the GlyGly-CTERM signal’s signature GG motif, and additionally showed that the cell’s type II secretion system (T2SS) is required for subsequent movement from the periplasm to the (correct) surface localization. In a parallel to PGF-CTERM proteins sorted by ArtA, GlyGly-CTERM proteins receive a new C-terminal attachment, in this case glycerophosphoethanolamine. The moiety is attached prior to interaction with the type II secretion system.

In additional bioinformatics work, we also described MYXO-CTERM, an orphan sorting signal because we were unable at the time assign a processing enzyme either homologous or analogous to the sortases(12), the exosortases and archaeosortases(7), or the rhombosortases(10). The MYXO-CTERM domain contains an invariant Cys residue in its signature motif, and often has two, close to each other but not adjacent. MYXO-CTERM appears on over 30 proteins in the deltaproteobacterial species *Myxococcus xanthus*, including the TraA protein later shown to be involved in the sharing of outer membrane proteins and lipids by compatible strains(13). As with rhombosortase substrates, MXYO-CTERM proteins likewise require processing by a T2SS system to reach the outer leaflet of the outer membrane(14).

Our continued efforts to expand the catalog of prokaryotic C-terminal sorting signals led to multiple new models, released over time in the TIGRFAMs(15) and the NCBIFAMs(16) collections of HMMs. MYXO-CTERM became, eventually, one of four orphan C-terminal sorting signals we defined that all share the property of featuring an invariant Cys residue in the signature motif. The similarities across these orphan sorting signals triggered further investigation, using phylogenetic profiling searches, examinations of conserved gene neighborhoods, and reasoning based on previously described patterns of design seen in prokaryotic protein-sorting systems(4,5,7,17). In this paper, we describe evidence that all four novel protein-sorting signals are recognized and processed by members of a different family of intramembrane proteases, related to the CAAX box-processing protease Rce1 (18) and its prokaryotic homologs (19,20).

## METHODS

### Identifying tripartite C-terminal sorting signals

We previously described several classes of prokaryotic C-terminally located protein-sorting signals of small size, and described the attributes typical of them that assist in their recognition (5,7,10). The signature attributes usually encountered include 1) location very close to the C-terminus, 2) multiple occurrences in a single genome, 3) a motif with at least three nearly invariant signature residues at the start of the homology domain, 4) a strongly hydrophobic region consistent with a transmembrane alpha helix, in the middle, 5) a cluster of basic amino acids, typically mostly arginine residues, two to five residues long, at the end. In addition, 6) most proteins sharing the sorting signal should have a recognizable signal peptide at the N-terminus, and 7) proteins sharing the sorting signal should include numerous pairs that lack regions of sequence similarity other than the sorting signal region itself. In many cases, 8) proteins with the sorting signal will have homologs from other lineages that either are shorter because the lack the signal, or that instead carry a different C-terminal sorting signal. In cases of *dedicated systems*, in which the relationship of sorting enzyme to target protein is one-to-one instead of one-to-many, regular co-occurrence of enzyme and target as products of consecutive or nearby genes may be observed instead of attributes 2, 7, and 8. Searches for novel classes of C-terminal protein-sorting signal were driven by curator-initiated investigations of select protein families or taxonomic clades, or by chance observations incidental to other protein family curation projects, rather than by programmatic search through all prokaryotic genomes.

### Identifying novel sorting enzyme families and variant forms

Multiple sequence alignments of known families of sorting enzymes were examined for clades with sufficient members to appear interesting, in which no matching sorting signal was yet described. Hidden Markov Models (HMMs), derived from curated multiple sequence alignments, were constructed and were given manually selected cutoffs and a name to use in RefSeq’s PGAP genome annotation pipeline(16). Whenever possible, HMMs for novel sorting enzyme variants, and for the cognate sorting signals, were built at the same time, with each family guiding the selection of proper cutoff scores for the other. To find entirely new classes of sorting enzyme, we searched by starting with orphan candidate sorting signals (those still without a known sorting enzyme) as the query, using Partial Phylogenetic Profiling(5) (see below), inspection for conserved gene neighborhoods, or both.

### Representative Genomes

From the set of over 14,000 representative complete and high-quality draft prokaryotic genomes, 6980 were selected randomly in June 2021. These *representative genomes* all were annotated by the Prokaryotic Genome Annotation Pipeline (PGAP) of the National Center for Biotechnology Information (NCBI) (16,21).

### Partial Phylogenetic Profiling

A diverse set of 5846 prokaryotic genomes (bacteria and archaea) from RefSeq was selected in July 2018 for use in Partial Phylogenetic Profiling (PPP) studies (the “*PPP genome set*”). The HMM for the MYXO-CTERM sorting signal, TIGR03901, was rebuilt, with 240 member sequences in the seed alignment, in November 2021. Sequences qualifying by HMM hit score were detected in 39 proteomes. Hits within twelve genomes of the order Myxococcales, within the class Deltaproteobacteria, numbered from 6 to 43. However, hits outside the Deltaproteobacteria all were singletons, several lacked the required Cys residue, most scored higher to different sorting signal HMMs (including NF033191 and NF038039), and all were judged to be false-positives. Three additional Myxococcales species, missed in the initial round of searching, were examined manually at this time, found each to have a sufficient number of valid although lower-scoring MYXO-CTERM domain-containing proteins, and were added to the phylogenetic profile. This gave a total of 15 curated true-positive genomes, out of 5846, to serve as a query profile for PPP.

The PPP algorithm has been described previously (5,22). It requires a phylogenetic profile to serve as query to use against the proteome of a selected genome. For each protein in the genome, PPP explores different possible sizes of protein family that the protein might be a part of. It looks at the fit between the list of species seen at a given family size and the query profile. The family size is varied by running down the list of top BLAST hits for the protein being evaluated and choosing an optimized stopping point where the score for the correspondence of species seen is the most unlikely to have been reached just by chance. The phylogenetic profiling is “partial” in the sense the score is based only on those species encountered in the collection of proteins examined in the BLAST hits list, which represents only a part of the full phylogenetic profile. The scoring system rewards hits to genomes marked as YES in the query profile, penalizes hits to genomes marked NO, but has no explicit penalty for YES genomes simply failing to show up in the BLAST hits list.

### SIMBAL analysis

Pfam (23) model PF02517 was used to identify CPBP family (*i*.*e*. Rce1-related) glutamic-type intramembrane proteases in the same proteomes as were used in Partial Phylogenetic Profiling. All members proteins from the 15 MYXO-CTERM true-positive proteomes were collected, yielding 87 proteins. These proteins became the YES set for SIMBAL (Sites Inferred by Metabolic Background Assertion Labeling) analysis (17,24). Searches from all other proteomes yielded 14390 proteins. No non-redundification was done.

### Clustering and phylogenetic trees of CPBP family proteases

Regions of 87 CPBP family proteases from 15 MYXO-CTERM-positive genomes were extracted with the aid HMM searches with PF02517. These domain sequences were aligned by MUSCLE(25). The alignment was visualized and trimmed in belvu (26). Clustering of the aligned sequences was performed by UPGMA (unweighted pair group method using arithmetic averages). Neighbor-Joining trees were generated in belvu, using the Storm and Sonnhammer distance correction method.

### Sequence Logos

Seed alignments for C-terminal protein-sorting signals were modified by removing alignment columns that had a gap character in more than half of sequences, and then the remaining sequences were made nonredundant by removal of sequences more than 80 % identical to others in the alignment. Sequence logos were built using the server at https://weblogo.berkeley.edu with default settings but custom coloring.

## Results

### Revising the MYXO-CTERM model

The model TIGRFAMs model TIGR03901, which has been described previously(13,15), was updated. The region modeled is short, about 34 amino acids, and highly divergent, so developing a broadly accurate is difficult. Optimizations that improve sensitivity and selectivity in one lineage tend to degrade performance in other lineage. A second version of the model was constructed with 240 sequences in the seed alignment, up from 123 in the first version. However, determining which proteins represent true members of the family requires curatorial review. Review established that MYXO-CTERM domains are restricted to two orders, Myxococcales and Bradymonadales, with the class Deltaproteobacteria. True-positive MYXO-CTERM domains occur close to the C-terminus, always contain a Cys residue in the signature motif region, and frequently contain two nearby Cys residues instead of just one. The sequence logo is shown in **Figure 1**. Accurate counting of MYXO-CTERM proteins in any one annotated genome requires an iterative process to build a lineage-specific custom model, as lineage-specific forms of the sorting signal and the sorting enzyme presumably co-evolve, and diverge from ancestral forms.

**Figure 1.**
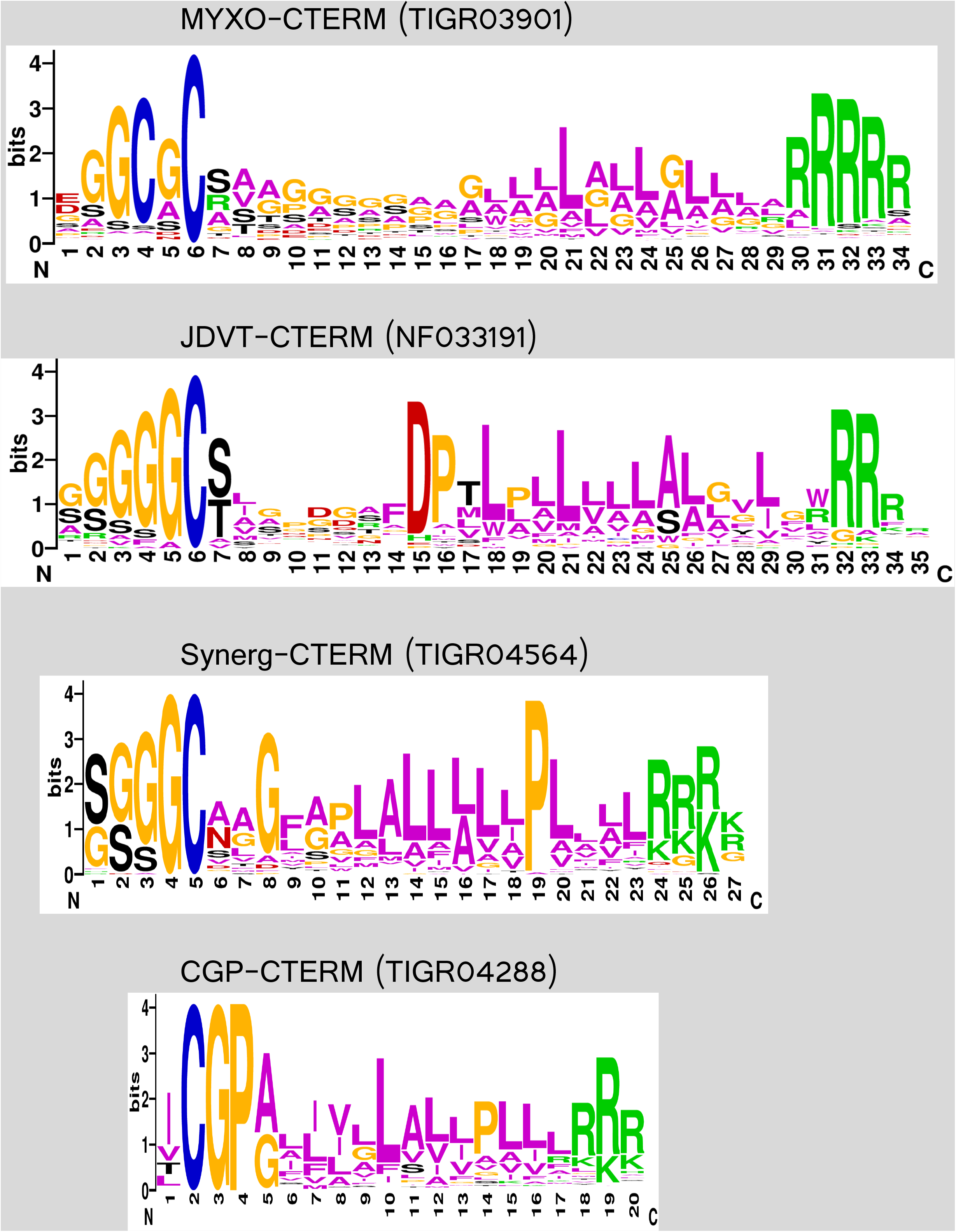
Sequence logos of Cys-containing tripartite C-terminal sorting signals. Sequence logos were built using https://weblogo.berkeley.edu, using default settings, but custom coloring to make Cys residues blue. Logos were constructed from seed alignments for their defining HMMs after removal of columns consisting mostl of the gap character. Seed alignment sequences come from non-overlapping sets of genomes. Logos are shown aligned on the invariant Cys residue of each domain. Myxo-CTERM (member proteins from the Myxococcales) is distinguished by an absence of charged or proline residues in the transmembrane (TM) helix region and frequent use of a second Cys residue (GCGC motif). JDVT-CTERM (member proteins from outside the Deltaproteobacteria) is distinguished by a single Cys only and a more Leu-rich TM segment preceded by a nearly invariant DP motif. Synerg-CTERM (member proteins from the Synergistetes) is shorter than MYXO-CTERM or JDVT-CTERM, with an invariant Pro in a leucine-rich stretch within the TM segment. The archaeal CGP-CTERM sorting signal, from the Thermococcales, is the shortest of all, with an invariant GP motif immediately following the invariant Cys.

### CGP-CTERM

We built HMM TIGR04288 (CGP-CTERM) originally for inclusion in the TIGRFAMs database(15), but have not previously described the domain in any publication. This putative protein-sorting domain occurs exclusively in and perhaps universally in *Thermococcus, Pyrococcus*, and *Palaeococcus*, the three genera of the order Thermococcales, all of which are hyperthermophilic archaea. Like MYXO-CTERM, the CGP-CTERM tripartite sorting signal domain contains a cysteine residue in its signature motif, which in this case is Cys-Gly-Pro. It is easily the shortest of the four Cys-containing sorting signals we describe here, just 20 amino acids in length.

Figure 1 shows the sequence logo for the CGP-CTERM, MYXO-CTERM, and two other novel sorting signals we describe here. The signature motif abuts the transmembrane segment with no spacer, as occurs for PEP-CTERM (cleaved by an exosortase), PGF-CTERM (cleaved by an archaeosortase), and GlyGly-CTERM (cleaved by rhombosortase), all of which are processed by deeply membrane-embedded enzymes.

Because the phylogenetic distribution of the CGP-CTERM domain is not sporadic at all, studies using the PPP algorithm are unlikely to be informative. A large number of proteins, 739, from *Pyrococcus horikos*hii OT3 all receive identical top scores from PPP, since a BLAST cutoff can be found each such that hits are registered for all 27 species with CGP-CTERM domains and for no species without. Because the apparent core proteome of CGP-CTERM domain-containing Thermococcales species is so large, PPP did not sufficiently narrow the search for the presumed sorting enzyme.

### Synerg-CTERM

Model TIGR04564 (Synerg-CTERM) likewise was built sufficiently long ago to include in releases of the TIGRFAMs database(15) before its move to the NCBI and inclusion within NCBIFAMs(16), but it too has never previously been described in a publication. The signature motif is a small Ser-rich and Gly-rich cluster that ends abruptly with a single invariant Cys residue. As with CGP-CTERM, there is no spacer between the signature motif and the transmembrane segment.

Sequences recognized by TIGR04564 occur so far in species such as *Dethiosulfovibrio peptidovorans, Aminiphilus circumscriptus, Aminomonas paucivorans, Fretibacterium fastidiosum, Cloacibacillus evryensis*, and *Synergistes jonesii*, but all of these belong to the order Synergistetes. Again, there appears not to be any extensive history of lateral gene transfer and gene loss, so phylogenetic methods would be expected to have limited utility. PPP was run for the proteome of *Dethiosulfovibrio peptidovorans* DSM 11002. The query profile has just seven genome assemblies. Twelve proteins receive top scores. Nine of these twelve belong to a cassette that encodes an apparently divergent subclass of type II secretion system (T2SS) operon. This strongly suggests that PPP is giving a meaningful signal, since sorting targets for both rhombosortase in *Vibrio cholerae*, and the presumptive myxosortase of *Myxococcus xanthus*, require a T2SS. The twelve protein list also includes and a glutamic-type intramembrane protease (WP_083797586.1), a member of the family described by Pfam model PF02517. This family includes the eukaryotic protease Rce1, which cleaves C-terminal CAAX box sorting signals after prior prenyl modification of the Cys side-chain(27), as well an archaeal protein capable of similar hydrolysis(19). Either or both of the two remaining proteins found by PPP, aspartate-semialdehyde dehydrogenase (WP_005660933.1), and a putative polysaccharide biosynthesis protein (WP_005659789.1), may not be directly relevant to protein sorting.

The Rce1 homolog co-occurring with Synerg-CTERM proteins is highly suggestive, since a membrane-embedded protease is exactly what is expected to process novel putative sorting signals. However, the genomes represented in the phylogenetic profile are fairly few and mutually rather closely related, so additional confirmation of the link between prokaryotic Cys-containing C-terminal sorting signals and Rce1 homologs is warranted.

### The JDVT-CTERM system

Efforts to improve the seed alignment and HMM used to detect MYXO-CTERM sequences led to identification of an apparently related sorting domain, differing in several key attributes. It occurs in a variety of Proteobacteria, including ***J****anthinobacterium* (Beta-proteobacteria), ***D****uganella* (Beta-proteobacteria), ***V****ibrio* (Gamma-proteobacteria), and ***T****hioalkalivibrio* (Gamma-proteobacteria), hence the name **JDVT**-CTERM. As **Fig. 1** shows, the tripartite architecture, presence of an invariant Cys residue, and overall length all resemble MXYO-CTERM. However, JDVT-CTERM has a nearly invariant Asp-Pro (DP) motif located nine residues C-terminal to the Cys, in the middle of the proposed membrane-spanning alpha-helix. It always has just one Cys residue, while MYXO-CTERM sorting signals frequently have two.

An even more profound difference from MXYO-CTERM systems is that true examples JDVT-CTERM are found typically just once per proteome, as computed for species that encode at least one such protein. This behavior suggests there should be numerous examples of a JDVT-CTERM domain-containing protein and its processing enzyme in the same operon, as seen with sortases and other protein-sorting enzymes that don’t have a general housekeeping role, but instead are dedicated to one target only. We took the top-scoring 38 examples of JDVT-CTERM proteins from a collection of representative genomes, made non-redundant to less than 80 percent pairwise identity, and then collected all proteins encoded with intergenic distances of 4000 nucleotides or less, and clustered them by performing a progressive alignment with Clustal-W(28). The largest single cluster, with 17 proteins, was a family of glutamic-type intramembrane proteases, relatively closely homologous to WP_083797586.1 that was putatively associated with the Synerg-CTERM system, and more distantly to eukaryotic type II CAAX prenyl-proteases(19,20,27). No other cluster contained more than 6 proteins. In 16 of 17 cases, the JDVT-CTERM and the intramembrane protease were adjacent, with no gene between them. This arrangement provides strong evidence of sorting target to sorting enzyme relationship.

Partial Phylogenetic Profiling (PPP) was performed, using RefSeq’s reannotation of *Thioalkalivibrio paradoxus* ARh 1 (GCF_000227685.2) as the model genome, and querying with the JDVI-CTERM profile (26 genomes out of 5846). The top score was achieved for WP_006748948.1, at a protein family size that reached 27 genomes total, 20 of them with JDVT-CTERM domain-containing proteins, for a score of 94.6. The next best score for any protein was 41.9 for WP_006747143.1, based on homologs found in 214 genomes, 19 of which had JDVT-CTERM domain-containing proteins. Because PPP scores are computed as the negative of the log of the odds of seeing such an extreme overrepresentation of YES genomes purely by chance, the results make it virtually certain that the co-occurrence of JDVT-CTERM domains and WP_006748948 family intramembrane proteases are connected by involvement in the same biological process. WP_006748948, another Rce1 homolog, shows convincing homology to the C-terminal half of the candidate sorting enzyme from the Synerg-CTERM system, WP_083797586.1, with the amino acid identity in that region exceeding 35 percent.

This second system analyzed, showing a links from sorting signal to an intramembrane protease both by gene neighborhood and by similar phylogenetic profiles, is exceptionally strong evidence for a direct biochemical relationship between a putative sorting enzyme and its JDVT-CTERM domain-containing targets.

### Partial Phylogenetic Profiling for the MYXO-CTERM system

Revision of the MYXO-CTERM model TIGR03901, construction of model NF033191 to describe the similar (but readily separable) JDVT-CTERM sorting signal, and manual review of questionable hits, typically one-per-genome hits outside the Deltaproteobacteria, made it possible to improve the phylogenetic profile used to represent the taxonomic range of the MYXO-CTERM domain. During this process, we identified the novel WGxxGxxG-CTERM domain, an orphan putative sorting signal, which occurs strictly outside the Deltaproteobacteria and which lacks the critical Cys residue. WGxxGxxG-CTERM is modeled by the NCBIFAMs HMM NF038039. Other than its utility to help judge the veracity of weak hits to model TIGR03901, it is not discussed further in this paper.

Following curatorial review, YES genomes in the MYXO-CTERM profile numbered 15, out of the 5846 in the PPP data set. Genome assemblies included GCF_000012685.1 (*Myxococcus xanthus* DK 1622) and GCF_001189295.1 (*Chondromyces crocatus*). PPP performed on *Chondromyces crocatus* returned 23 proteins with perfect scores, all 15 YES genomes found when BLAST cutoffs reach exactly 15 genomes. One of these, WP_082362253.1, is an intramembrane protease with a C-terminal region homologous to WP_006748948.1 of the JDVT-CTERM system and to the C-terminal half of WP_083797586.1 from the Synerg-CTERM system. These findings strongly suggest that WP_082362253.1 (MXAN_2755) is the previously cryptic myxosortase for MYXO-CTERM proteins in *Myxococcus xanthus*. We rename this protein MrtX, that is, a **m**yxoso**rt**ase of the type seen in *M*. ***x****anthus*.

### Clustering and Phylogeny of Rce1 homologs in MYXO-CTERM system genomes

To address the question of whether multiple mysosortases might share responsibilities for recognizing and cleaving MYXO-CTERM sorting signals, we collected all 87 members of family PF02517 from our 15 curated MYXO-CTERM-positive species. A multiple sequence alignment showed a core homology region, lining up well with the homology domain described by PF02517. The alignment of the core region is shown in **Figure 2**. Aligning full-length sequences (not shown), then sorting the sequences by average percent identity as computed within the resulting untrimmed multiple sequence alignment, revealed a number of different clusters with no more than one member per species represented and with higher levels of sequence identity in the core region than is ever seen between two different paralogs from a single. Only one cluster had representatives from all 15 species. The two paralogs of MrtX in *Myxococcus xanthus* belonged to the next two largest clusters, WP_011553332.1 in an eight member cluster, and WP_011555198.1 in a six-member cluster.

**Figure 2.**
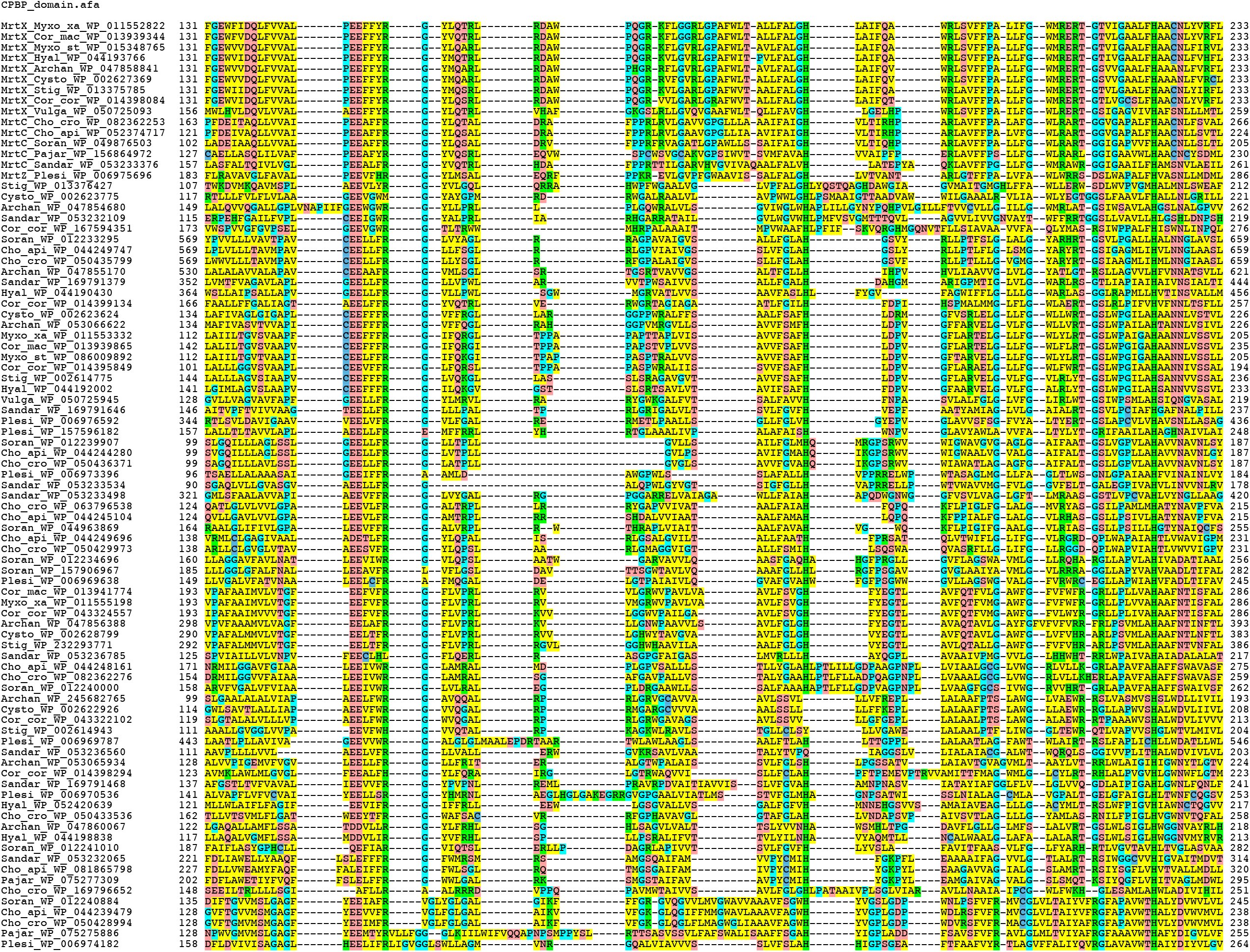
Sequence alignment of CAAX prenyl-protease homology domains. Homologous regions were excerpted from all 85 members of Pfam domain family PF02517 found in any of the 15 genomes identified in the set of MXYO-CTERM-positive proteomes used for PPP analysis. In the color scheme used, yellow (V,I,L,M,A,W,F,Y) indicates hydrophobic, green (R,K,H) indicates basic, red (D,E,Q,N,S,T) indicates acidic or neutral but hydrophilic, light blue (G, P) indicates residues common in turns, and dark blue indicates C (rare and frequently involved in disulfide bond formation, metal-binding, or catalysis). Sequences are grouped hierarchically by amino acid percent identity, using UPGMA (unweighted pair group method using arithmetic averaging), with the 15 sequences of the myxosortase cluster at the top. The EE motif at positions 31-32 contains the primary catalytic site.

The UPGMA tree for the untrimmed alignment is not shown, as clustering by percent identity does not show phylogenetic relationships, and additional domains N-terminal or C-terminal to the core homology domain could mislead. Instead, we show a neighbor-joining tree, computed from a newly constructed alignment of just the core homology domain, visualized using FigTree version 1.4.4 (http://github.com/rambaut/figtree/) (see **Figure 3**). *M. xanthus* protein MrtX (WP_011552822) belongs to the only cluster with members from all 15 species, while paralogs WP_011553332.1 and WP_011555198.1 belong to the two next largest clusters, with sizes of eight and six, respectively. The 15-member cluster is notable because member sequences have such high levels of sequence identity in the protease domain region, above 40 % identity even between the most distant pairs, while sequence similarity is barely detectable between two very different types of N-terminal domain. In the region of homology shared by all Rce1 homologs in MYXO-CTERM-positive species, the 15-member cluster is not only the one cluster with a member from every required species, nearly twice the size of the next largest cluster. It is also the mostly highly conserved cluster.

**Figure 3.**
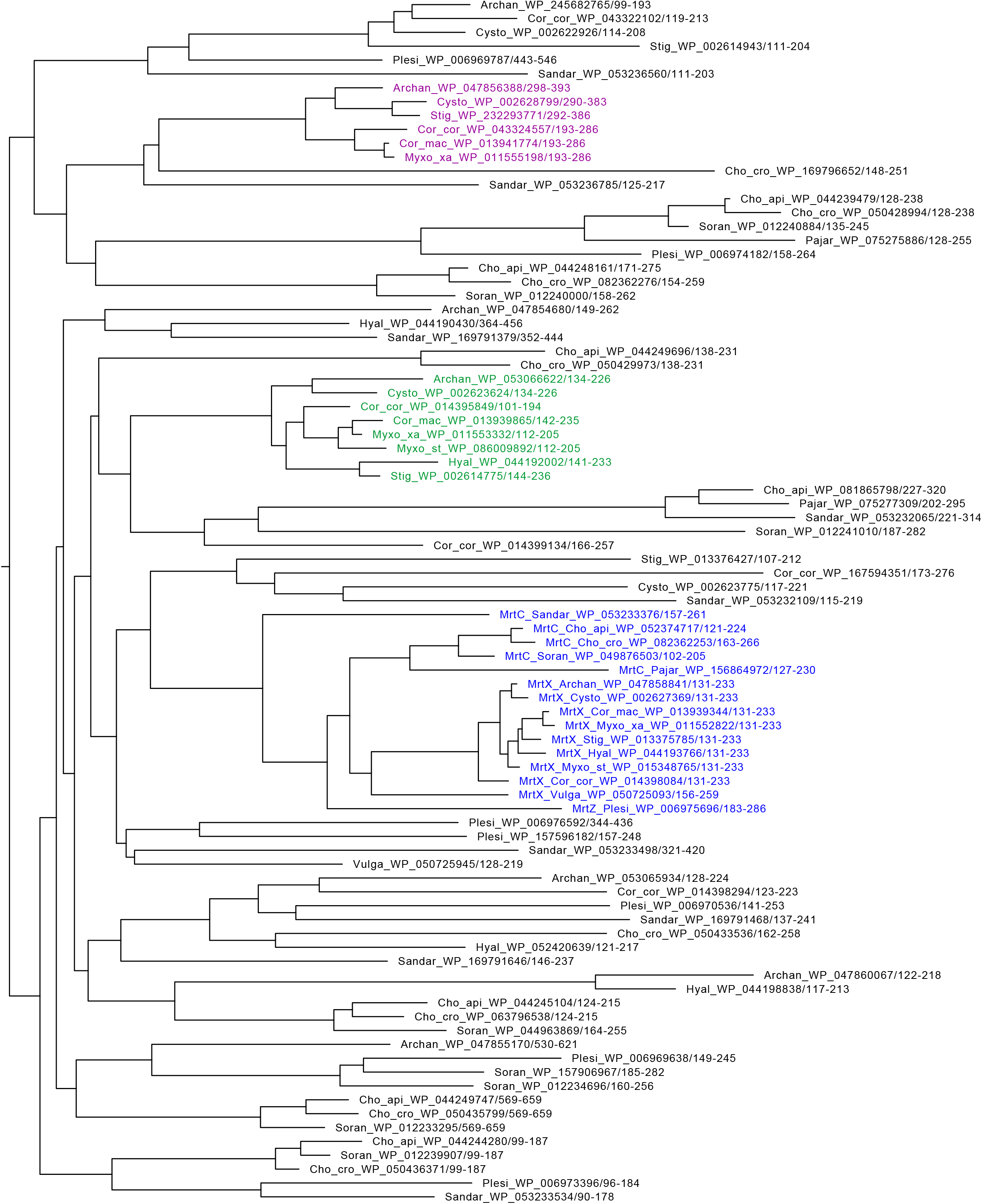
Neighbor-joining (NJ) tree of all CAAX prenyl-protease homology domains from fifteen bacteria with MXYO-CTERM protein-sorting systems. The NJ tree was constructed from the multiple sequence alignment shown in Figure 2, using Storm and Sonnhammer distance correction in belvu(26). The tree was exported to FigTree v.1.4.4 for display. The tree is unrooted, and is shown with horizontal terminal branches for legibility. The cluster containing all myxosortases, such as MrtX from *Myxococcus xanthus*, with 15 members, is shown in blue. The two clusters that have non-myxosortase paralogs from *M. xanthus* are colored green (with 8 members) and purple (with 6 members). Species abbreviations that prefix the RefSeq protein accession numbers (starting “WP_”) are Archan (*Archangium gephyra*), Cho_api (*Chondromyces apiculatus*), Cho_cro (*Chondromyces crocatus*), Cor_cor (*Corallococcus coralloides*), Cor_mac (*Corallococcus macrosporus*), Cysto (*Cystobacter fuscus*), Hyal (*Hyalangium minutum*), Myxo_xa (*Myxococcus xanthus*), Myxo_st (*Myxococcus stipitatus*), Pajar (*Pajaroellobacter abortibovis*), Plesi (*Plesiocystis pacifica*), Sandar (*Sandaracinus amylolyticus*), Soran (*Sorangium cellulosum*), Stig (*Stigmatella aurantiaca*), and Vulga (*Vulgatibacter incomptus*).

**Figure 4.**
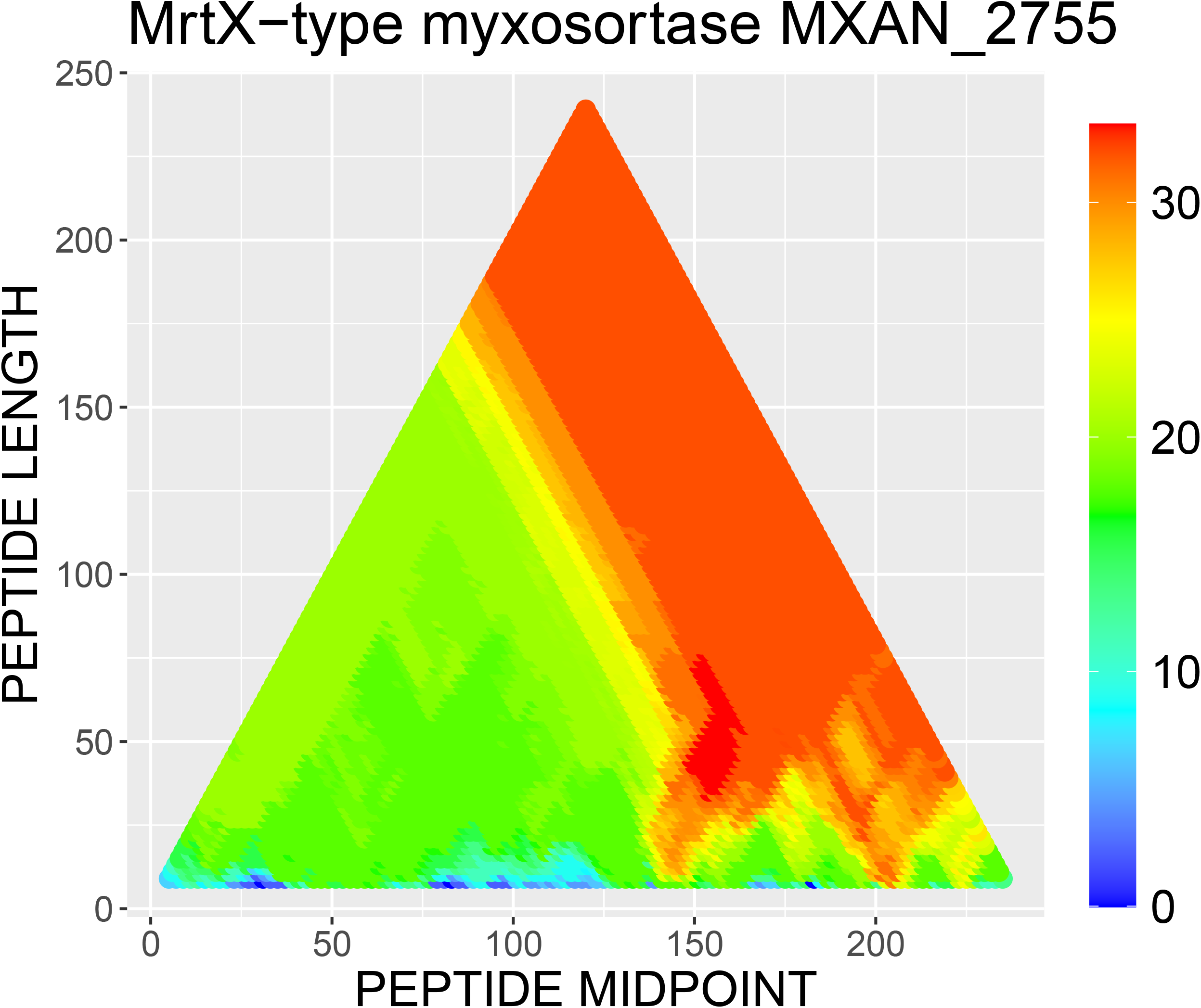
SIMBAL plot of MrtX from *Myxococcus xanthus* (MXAN_2755). The training set’s YES partition consisted of 85 proteins identified by Pfam model PF02517 in the 15 proteomes identified as having MYXO-CTERM sorting signals in collection used for Partial Phylogenetic Profiling. The NO partition consisted of PF02517 from all other species. Proteins in the YES partition were 0.6% of all proteins. Red indicates high SIMBAL scores, that is, an exceptionally strong skew toward the YES set. The protein length is 239, so center points for the minimum size window tested, 9 amino acids, occur at positions 5 through 235. For peptide lengths shorter than 50, scores for peptides centered in the N-terminal half of the protein score no higher than 20 (green), based on BLAST hits to 9 YES proteins and 0 NO proteins, as only 9 of 15 species have an MrtX protein with its distinctive N-terminal domain. Notably high scores occur for peptides of length 13 (near the bottom of the figure) centered around 147 and around 203. These stretches of sequence, 141-VVALP**EE**FFYRGY-153 (glutamic acid residues shown in boldface) and 197-LSVFFPALIFGWM-209, have SIMBAL scores of 29.0 (14 YES vs. 2 NO) and 31.1 (14 YES and 0 NO). These locally high scores suggest the two regions contain residues particularly important for conferring specificity for myxosortase activity among members of the broader family of glutamic-type intramembrane proteases.

### SIMBAL analysis of the *Myxococcus xanthus* myxosortase enzyme

The clear identification of WP_006748948.1 as a protein-sorting enzyme for the JDVT-CTERM system, and of additional members of Pfam family PF02517 as probably sorting enzymes for similar Cys-containing sorting signals, raised an important question. Do multiple enzymes Rce1-like paralogs in single proteome share in the processing of the target proteins with the tripartite sorting signals described in this paper? Or is it more typical that just one family enzyme is the housekeeping myxosortase, while other members of family PF02517 have very different responsibilities?

*Chondromyces crocatus* assembly GCF_001189295.1 has nine members of family PF02517. These are WP_082362253.1(renamed myxosortase C, or MrtC), WP_050435799.1, WP_063796538.1, WP_050429973.1, WP_050433536.1, WP_050436371.1, WP_169796652.1, WP_082362276.1, WP_050428994.1). Myxococcus xanthus has three paralogs, namely WP_011552822.1 (myxosortase X, or MrtX), WP_011553332.1, and WP_011555198.1. The strongest pairwise match among any of these twelve proteins is between MrtC and MrtX, with that similarity apparently restricted to the C-terminal portions of the two proteins, as the N-terminal domains appear unrelated. Amino acid sequence identity exceeds 50% in the shared C-terminal domain. The proposed sorting enzyme of the Synerg-CTERM proteins, WP_083797586.1, is more closely related to MrtC and MrtX than to any of their paralogs.

Similarly, the sorting enzyme WP_006748948.1 of the JDVT-CTERM system, here renamed MrtJ (myxosortase-like sorting protein of the JDVT-CTERM system) is more closely related to MrtX than to its paralogs. The results of all these comparisons makes it seem likely that a single mxysortase enzyme, not a group of several paralogs, handles the processing of MYXO-CTERM-like sorting signals in each of the species we examined.

SIMBAL analysis was performed using a sliding window 17 amino acids long. All three paralogs from the glutamic-type intramembrane protease were tested. For the paralog WP_011553332, no score for any substring scored better than 18.99, hitting 9 proteins from the YES partition, 1 from the NO partition, while only 0.6% of proteins are in the YES set. For paralog WP_011555198, the top scores were for 17 from the YES set, 182 from the NO set (scoring 13.98), or 6 from the YES set, 0 from the NO set (scoring 13.33). In contrast, SIMBAL analysis for the actual myxosortase MrtX, using BLAST searches for sequence regions as short as 13 amino acids long, produced SIMBAL scores as high as 26 (14 YES vs. 2 NO) for the sequence centered at 147, 31 (14 YES and 0 NO) when centered at or near 203. Slightly longer sequences, and very long or full-length sequence produce top scores, either 32.11 for 15 YES vs. 1 NO, or 32.31 (15 YES vs. 0 NO). The SIMBAL data point for the full length sequence effectively reproduces the result from Partial Phylogenetic Profiling, as that analysis is based on full-length sequences only.

### Expanding the Set of MYXO-CTERM-positive species and Myxosortase subtypes

Myxosortase activity may be found in a single homology region shared by different types of myxosortases found in different lineages. We built a new HMM, NF040914, to describe the conserved region. SIMBAL analysis consistently showed maximum scores with 15 true-positive hits and 1 apparent false-positive. A search of the negative set with NF040914 identified a single protein scoring above cutoff, WP_012632673.1, from *Anaeromyxobacter dehalogenans*. Repeated the search for overlooked MXYO-CTERM proteins in the species identified just two, both low-scoring vs. TIGR03901, namely WP_150106367.1 and WP_015934593.1. Examination showed sorting signal sequences of SGGCGAGGTGALAMIGAAALAALRRRKP and AVGCQAGAGSGWALLAPLAVVAAAALRRRRQR, both of which we judge to be true MYXO-CTERM sequences. We elect not correct profiles and training sets after this determination, however, as could introduce curator bias to a statistical analysis.

A search of a set of over 14000 bacterial proteomes identified as “representative” by NCBI in May of 2022, with the myxosortase core domain model (NF040914), identified five species outside the Myxococcales. All five (WP_111331193.1, WP_141199628.1, WP_146979376.1, WP_111728004.1, and WP_127778827.1) are from another Deltaproteobacterial lineage, the Bradymonadales. Their five species (*Bradymonas sediminis, Persicimonas caeni, Lujinxingia vulgaris, Lujinxingia litoralis*, and *Lujinxingia sediminis*) have abundant, easily detected MYXO-CTERM, validating their identity. However, their overall domain architecture differs, with the inclusion of an additional central domain about 100 amino acids in length. The myxosortases of the Bradymonadales are designated MrtP (NF040674).

## Discussion

Myxosortase previously has been hard to identify in the proteomes of the taxonomic order Myxococcales (and the related Bradymonadales) because it is just one of large number of protein families well-conserved in those proteomes and absent outside. Worse, its architecture is variable, so the N-terminal halves of myxosortases from two different species may be unrelated. Because a single myxosortase acts as a single copy housekeeping enzyme, with responsibility for processing different target proteins that require expression at different time, myxosortase is likely not co-regulated with any one MYXO-CTERM protein. It is therefore not encoded in the same operon as its target proteins, and could not be discovered by shared gene neighborhood. MYXO-CTERM therefore remained an orphan sorting signal, with the responsible processing enzyme remaining unknown.

A further complication was the existence of other sorting signals that also carry an invariant Cys residue, are similarly small in size, and were therefore non-trivial to separate. In fact, it was the effort to deconvolute true MYXO-CTERM sequences from others that score similarly in HMM search results that led to our first detection of the JDVT-CTERM sorting system. Because that system, found strictly outside of the Myxococcales, is a *dedicated system*, with a single protein target per enzyme, co-regulated and co-operonic, clues from conserved gene neighborhood supplemented co-occurrence evidence detected by PPP.

The variable architecture we see for Rce1 homologs in the bacteria we examined here suggests that many, including myxosortases MrtX, MrtC, and MrtP, may be bifunctional. That is, two separate modifications may occur, perhaps first lipid attachment to one or more Cys residues, then cleavage distal to the most C-terminal Cys residue. This matches the model proposed by Sah, et al.(14), who provided experimental evidence that cleavage occurs, a Cys residue in the MYXO-CTERM region is required, multiple MYXO-CTERM proteins become exposed on the cell surface, and a type II secretion system is required for MYXO-CTERM proteins to reach their surface destinations.

This work unites at least three small, prokaryotic, C-terminally located, membrane-spanning, Cys-containing protein-sorting signals, all bacterial, as sharing the same class of previously unrecognized sorting enzyme – a class familiar to many because some eukaryotic members of the family act on CAAX box proteins in the endoplasmic reticulum. A fourth tripartite-pattern sorting signal, the archaeal CGP-CTERM domain (TIGR04288), is suspected to be handled in a similar way, although that has not yet been shown conclusively.

As more novel genomes are sequenced, and the power of comparative genomics-driven approaches continues to grow, additional novel sorting systems are likely to be discovered. The WGxxGxxG-CTERM domain, for example, is now detected by HMM model NF038039 and is seen to be broadly distributed, with the pattern of design of a tripartite sorting signal and the familiar many-or-none distribution familiar from most LPXTG, PEP-CTERM, and MXYO-CTERM-containing species. It is therefore a putative novel sorting signal, but it remains an orphan sorting signal for now.

## ACKNOWLEDGEMENTS

This research was supported by the National Center for Biotechnology Information of the National Library of Medicine (NLM), National Institutes of Health.

